# Effects of climatic change on the potential distribution of *Lycoriella* species (Diptera: Sciaridae) of economic importance

**DOI:** 10.1101/2021.07.23.453546

**Authors:** Roberta Marques, Juliano Lessa Pinto Duarte, Adriane da Fonseca Duarte, Rodrigo Ferreira Krüger, Uemmerson Silva da Cunha, Luís Osorio-Olvera, Rusby Guadalupe Contreras-Díaz, Daniel Jiménez-García

## Abstract

*Lycoriella* species (Sciaridae) are responsible for significant economic losses in greenhouse production (e.g. mushrooms, strawberry, and nurseries). Current distributions of species in the genus are restricted to cold-climate countries. Three species of *Lycoriella* are of particular economic concern in view of their ability to invade across the Northern Hemisphere. We used ecological niche models to determine the potential for range expansion under climate change future scenarios (RCP 4.5 and RCP 8.5) in distributions of these species of *Lycoriella*. Stable suitability under climate change was a dominant theme in these species; however, potential range increases were noted for key countries (e.g. USA, Brazil, and China). Our results illustrate the potential for range expansion in these species in the Southern Hemisphere, including some of the highest greenhouse production areas in the world.

## 1. Introduction

Sciaridae (Insecta, Diptera), known as black fungus gnats, comprise more than 2600 species worldwide, most of which are harmless to human activities [1]. Although most of the species have phyto-saprophagous larvae, 10 known species have larvae that may feed on living tissue, damaging roots or mining stems and leaves of economically important crops and ornamental plants, which can lead to significant economic losses [2–5].

Mushroom crops can be affected severely by sciarids. Sciarid larvae can feed on the developing mycelium inside the substrate and destroy sporophore primordia. Mature mushrooms may also be damaged by larvae tunneling into the tissue, which leads to product depreciation. Severe larval infestations may even destroy the sporophores, causing severe economic losses to producers [6].

Since 1978 worldwide production of cultivated edible fungi has increased around 30-fold and is expected to increase further in coming years [7]. Mushrooms represented a global market of US$63B in 2013 [8]. According to the USDA, the value of mushroom sales for 2019-2020 in the USA was US$1.15B, up 3% from the previous season [9]. Among the mushrooms produced, *Agaricus bisporus* is the most important, according to the Economics, Statistics and Market Information System. In 2020-2021, the area under production is 12,470 m^2^, 56.5% of which is in Pennsylvania territory [9].

The mushroom industry has suffered major economic losses caused by sciarid larvae in Australia, USA, Russia, United Kingdom, and South Korea [10,11]. Three sciarid species of the genus *Lycoriella* Frey, 1942 (*L. agraria, L. ingenua*, and *L. sativae*) are particularly harmful to cultivated mushroom crops, and are considered to rank among the most important pests of cultivated mushrooms throughout the world [4,10]. In countries like the United States and England, *L. ingenua* and *L. sativae* are the most serious pests in mushroom crops [12], as well in Europe [10]. In Korea, *L. ingenua* is considered as the most economically important [11]. Given their small size, sciarid larvae can be transported inadvertently to new areas by human activities. Infested potting mix, soilless media, commercial plant substrate, and rooted plant plugs have been shown to act as pathways for sciarid movement [13]. From 1950 onwards, globalization promoted transporting these invasive species [14]. In this sense, studies of their ecology, environmental requirements, and climatic change impacts for establishment of invasive populations are needed.

Ecological niche modeling (ENM) is used to evaluate relationships between environmental conditions and species’ abundances and occurrences [15]. Understanding potential distributions of species represents an important opportunity for pest control and mitigation of possible invasors (e.g. Compton et al., 2010; Gallien et al., 2010; Thuiller et al., 2005). Considering that the three *Lycoriella* species are economically important and are invasive species [10,19], niche modeling allows researchers to identify areas not currently occupied by them; if dispersal is possible or facilitated, these areas can be invaded and populations established in these regions [15]. For these reasons, we used ENM to identify new regions of potential invasive risk for three *Lycoriella* species with pest status in mushroom production, under current and future climate conditions (2050) for two greenhouse gas emissions scenarios.

## 2. Materials and Methods

### 2.1 Occurrence data

Occurrence data for *Lycoriella* species were obtained from published papers available in bibliographic databases (Google Scholar, Web of Science, Scopus), and from SpeciesLink (http://splink.cria.org.br/) and GBIF (http://www.gbif.org). We gathered all data from 1950-2018 for synonyms [3] including *L. agraria* [20] and its synonym *Sciara multiseta* [21], *L. ingenua* [22] and its synonym *S. pauciseta* [23] and *L. sativae* [24], and its synonyms *L. auripila* [25] and *L. castanescens* [26]. Occurrences lacking geographic coordinates were georeferenced in Google Earth (2015; https://earth.google.com/web/). We excluded records lacking the exact location or with high geographic uncertainty (e.g. name of the country as a collection site).

We assembled the occurrence data for each *Lycoriella* species, and performed a geographic spatial thinning such that no thick points were closer than 50 km using the spThin R package [27]. As such, we used 43 *L. agraria* occurrences, 118 *L. ingenua* occurrences, and 136 *L. sativae* occurrences. Finally, the data were split randomly into two subsets: 50% for model training and 50% for model testing (Suppl. information figures 1, 2 and 3).

**Figure 1.**
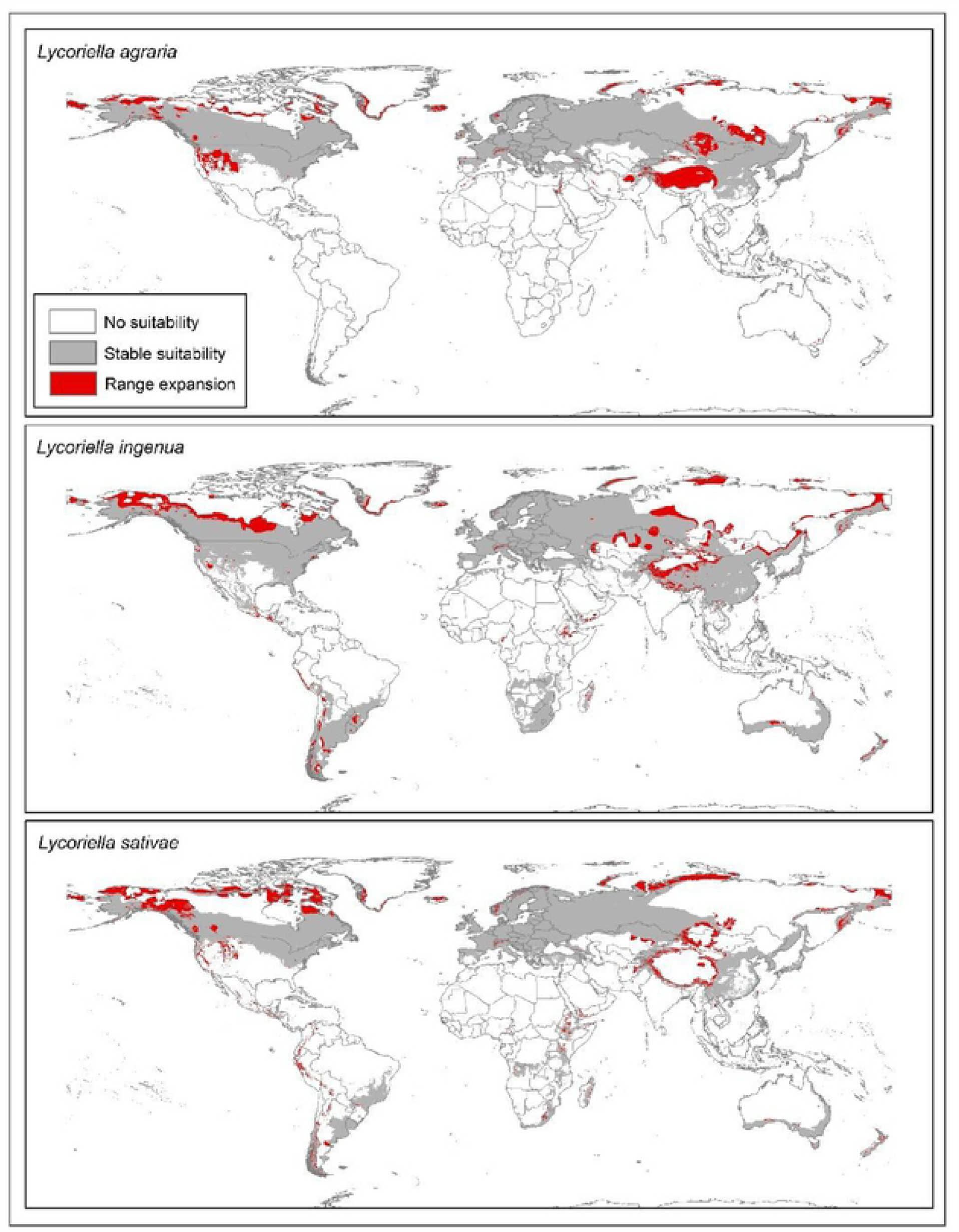
Potential distributions of three *Lycoriella* species under present and future climate conditions under two emissions scenarios (RCP 4.5 and RCP 8.5).Models show potential for range expansion worldwide in areas with low extrapolation risk.

**Figure 2.**
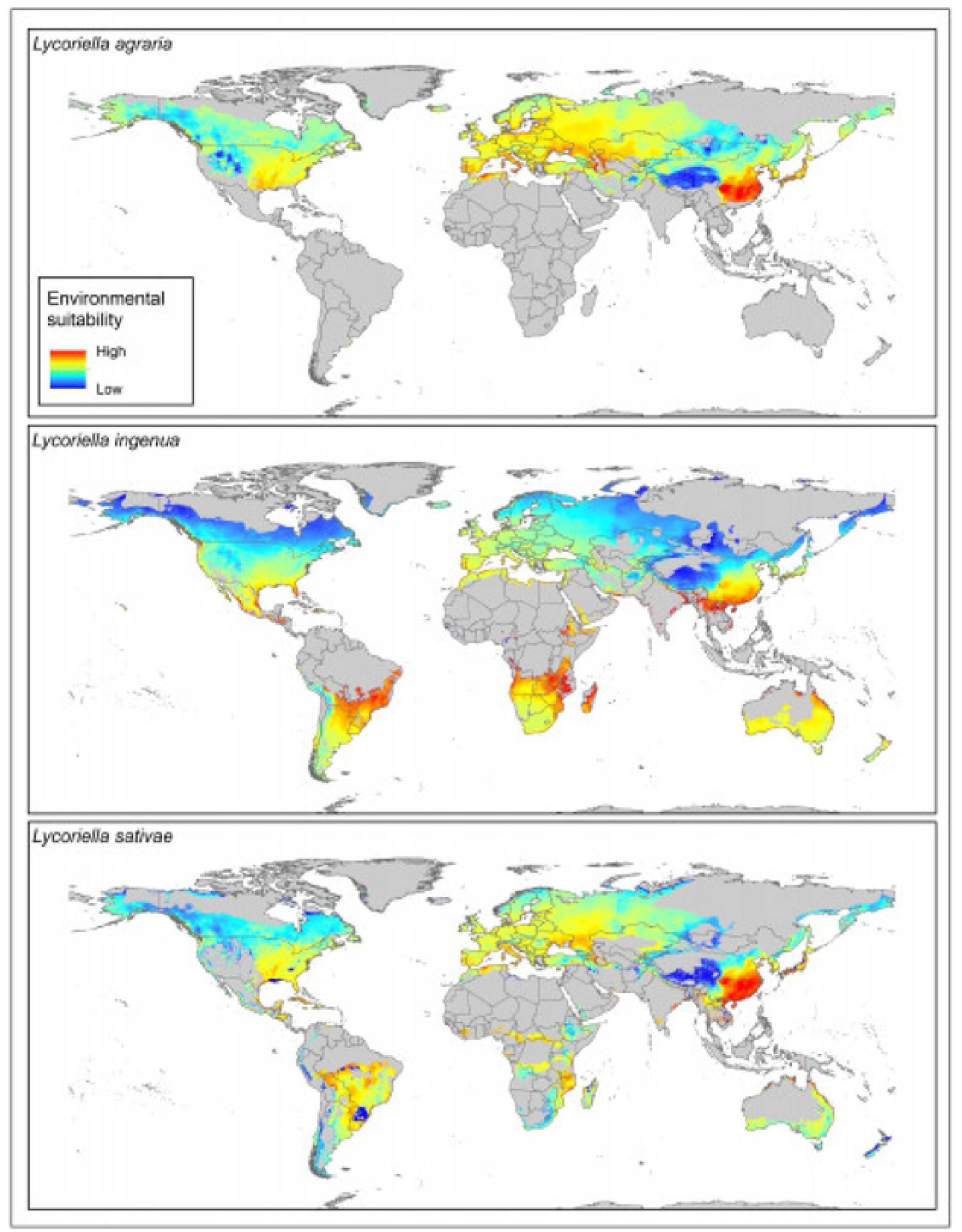
Environmental suitability for three *Lycoriella* species under current climate conditions worldwide.

### 2.2 Environmental variables

The bioclimatic variables used here to summarize climatic variation were from WorldClim version 1.4 [28]; we excluded four variables (bio 8, bio 9, bio 18, bio 19) that present spatial artefacts [29]. We summarized future conditions via 22 general circulation models (GCMs; Suppl. information figures 4, 5 and 6) for 2050 available from Climate Change, Agriculture and Food Security [30]. Two greenhouse gas emissions scenarios (RCP 4.5 and RCP 8.5) were used to explore variation among possible future emissions trajectories. The climate variables were used at a spatial resolution of 2.5 min (∼5 km^2^). We used Pearson’s correlations across each of the calibration areas for each species, removing one from each pair of variables with correlation *≥*0.80. The remaining not correlated variables were grouped into all possible sets of ≥2 variables for testing (Cobos et al., 2019; Table 1).

**Table 1.**
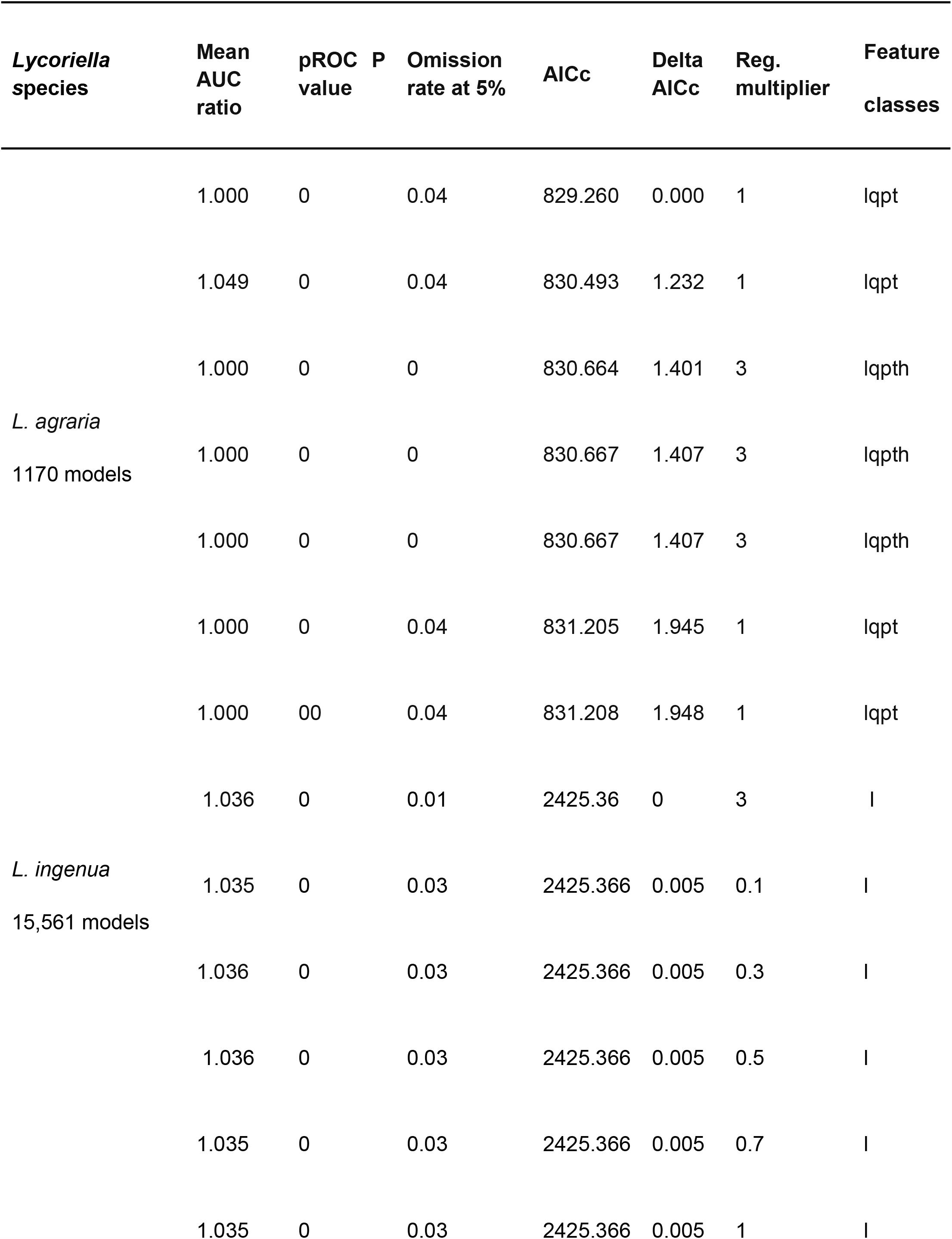
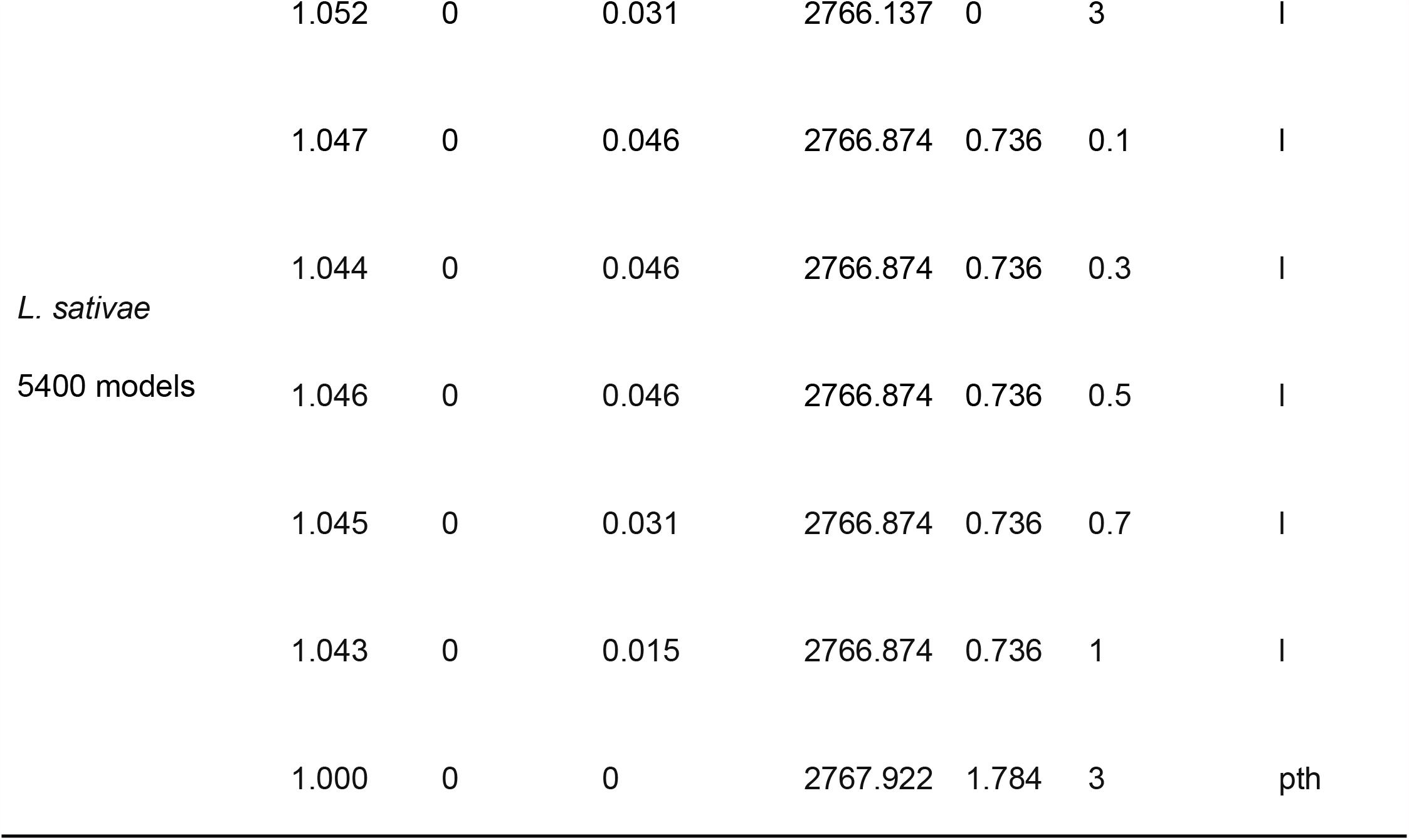
Best models selected and evaluated based on statistical significance (partial ROC), performance (omission rates: OR), and complexity (AICc). This model was calibrated and projected using the environmental variables shown in Table 2.

### 2.3 Model calibration and evaluation

We calibrated candidate models in Maxent 3.4.1 (Phillips et al., 2006), and model selection was achieved using the kuenm R package [32]. We assessed all potential combinations of linear (l), quadratic (q), product (p), threshold (t), and hinge (h) feature types; in tandem with 9 regularization multiplier values (0.1, 0.3, 0.5, 0.7, 1, 3, 5, 7 and 10); and the 26, 247, and 120 environmental data sets described above, for *L. agraria, L. ingenua*, and *L. sativae*, respectively. We therefore explored 1170 candidate models for *L. agraria*, 15,561 for *L. ingenua*, and 5400 for *L. sativae* (Table 1). We evaluated significance, performance, and complexity, of each candidate model, to choose optimal parameter settings, as follows. Significance testing was via partial receiver operating characteristic (pROC) tests [33]; values of partial ROC were calculated based on maximum acceptable omission error rate of *E* = 0.05. Omission rates were determined using a random 50% of the occurrence data, and model predictions were binarized via a modified least training presence thresholding approach (*E* = 0.05). Finally, we evaluated model complexity using the Akaike information criterion with correction for small sample size (AICc), following Warren and Seifert (2011). All modeling processes were included in the kuenm R package [32].

We use a hypothesis of the accessible area (**M**) for each species to calibrate our models [35,36], using buffers of 50 km around occurrence data points remaining after spatial thinning. Final models were taken as the median of the 10 replicates for best models and were projected worldwide. Model summaries were generated from thresholded median model projections (Figure 2) using the *E* = 0.05 value. We used the kuenm package [32] for these final steps as well. For each future-climate scenario (RCP 4.5 and RCP 8.5), we transferred the models and evaluated extrapolation conditions through MOP analysis [37], using the ntbox R package [38].

We summarized the projections of the models as medians of the replicate models using a modified least presences threshold value of *E* = 0.05. Binary maps for future conditions were used to determine uncertainty in terms of disagreement among predictions from the different GCMs (Suppl. information figures 4, 5 and 6). We summed the maps and used overlap between present and future potential distribution areas to determine prediction stability and range increase for each species in geographic areas with low extrapolation risk based in MOP analysis (Supp. information figures 7, 8 and 9).

## 3. Results

We created and evaluated 22,131 candidate models for the three *Lycoriella* species, (Table 1). For *L. agraria*, of 1170 candidate models, 669 were significant (*P < 0*.*05*) and 575 had omission rates below 5%; of significant, low omission models, 7 were selected according to low complexity (AICc; Table 1). Of 15,561 candidate models for *L. ingenua*, 6898 were significant and 6789 models had omission rates below 5%; we selected 6 models based on complexity. Finally, we generated 5400 candidate models for *L. sativae*, of which 1323 were significant and 1061 had omission rates below 5%; we selected 7 models according to AIC criteria (Table 1).

Nine variables were identified as key in our ENMs (Table 2). In general, *Lycoriella* species showed relationships with seasonality in temperature and precipitation, and with variables related to cold temperatures and wet seasons (Table 2), with variable contributions ranging 4.6-49.8%. The maximum number of variables for best models was in *L. sativae*, including high differences in variable contribution (Table 2).

**Table 2.**
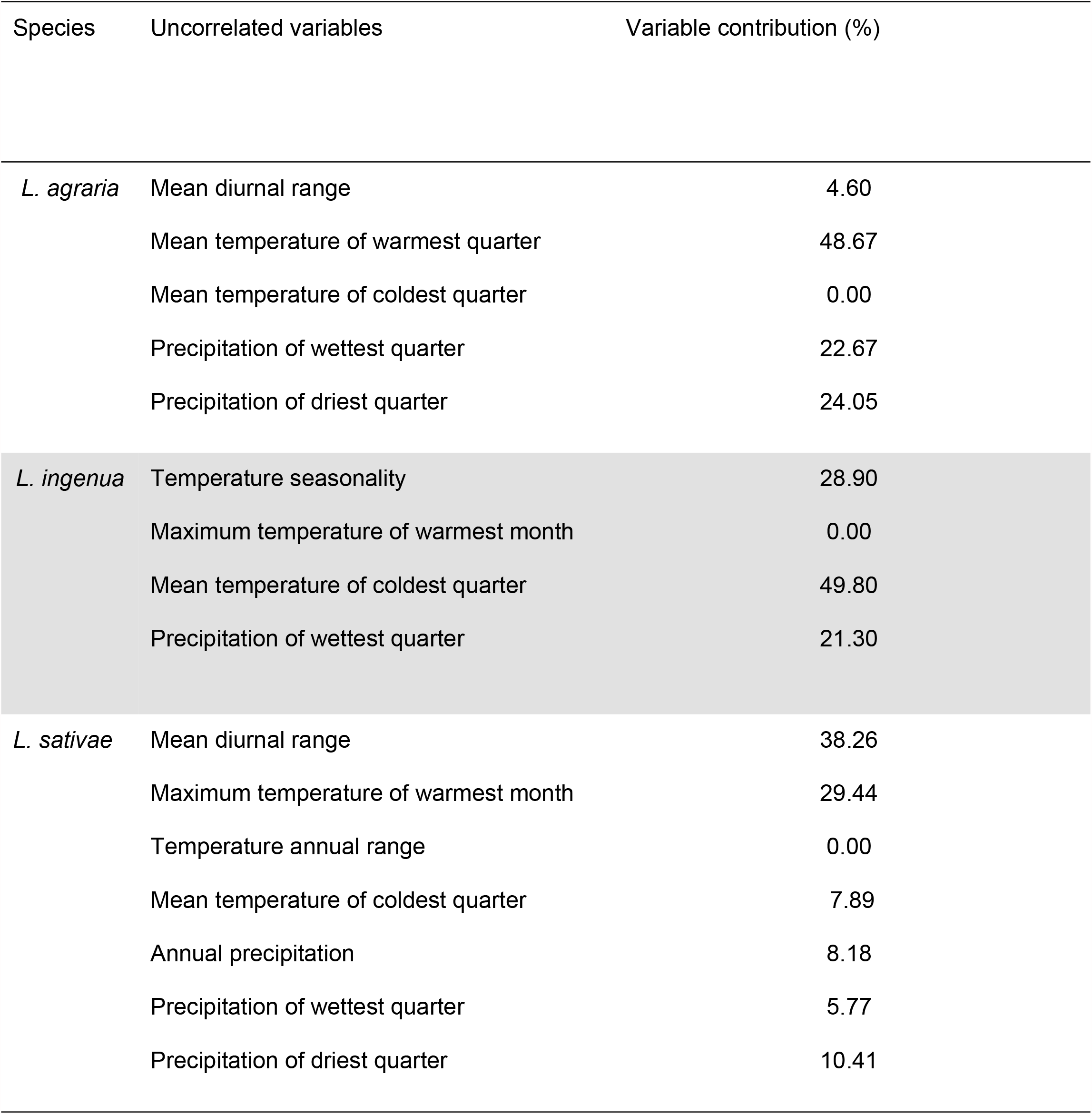
Models and variables that were relatively uncorrelated (Pearson’s correlation ≤ 0.8) for *Lycoriella* species. The models were built and tested used 26 variables sets for *L*.*agraria*, 247 variables sets for *L. ingenua*, and 120 variables sets for *L. sativae*.

| Species | Uncorrelated variables | Variable contribution (%) |
| --- | --- | --- |
| *L. agraria* | Mean diurnal range | 4.60 |
|  | Mean temperature of warmest quarter | 48.67 |
|  | Mean temperature of coldest quarter | 0.00 |
|  | Precipitation of wettest quarter | 22.67 |
|  | Precipitation of driest quarter | 24.05 |
| *L. ingenua* | Temperature seasonality | 28.90 |
|  | Maximum temperature of warmest month | 0.00 |
|  | Mean temperature of coldest quarter | 49.80 |
|  | Precipitation of wettest quarter | 21.30 |
| *L. sativae* | Mean diurnal range | 38.26 |
|  | Maximum temperature of warmest month | 29.44 |
|  | Temperature annual range | 0.00 |
|  | Mean temperature of coldest quarter | 7.89 |
|  | Annual precipitation | 8.18 |
|  | Precipitation of wettest quarter | 5.77 |
|  | Precipitation of driest quarter | 10.41 |

Current suitable areas for *Lycoriella* species includes much of the Northern Hemisphere, except for parts of Greenland, Russia, and northern China. *L. ingenua* and *L. sativae* also had suitable areas in the Southern Hemisphere: South America, southern Africa, and Australia (Figures 1 and 2). The model for *L. agraria* indicated high suitability in parts of North America, except Mexico (Figures 1 and 2), as well as much of Eurasia except for Russia, the Indian Subcontinent, and Southeast Asia. Suitable areas for *L. ingenua* were indicated for much of the Americas, except for parts of Canada, Alaska, Central America, and northern South America. *Lycoriella sativae* showed high suitability in the Americas, except in the western United States, northern Canada, central Mexico, and parts of South America (e.g. northern Brazil, Pacific Coast). Eastern and southern Asia was not suitable for this species; nor were much of Australia, North Africa, or parts of central and southern Africa.

Stable suitable conditions for the three *Lycoriella* species were the dominant pattern in comparisons of current and future potential distributions (Figure 1 and suppl. information figures 4, 5 and 6). Potential range expansion for the three species were noted in North America and Southeast Asia (Figure 1 and suppl. information figures 4, 5 and 6). Range reductions were detected in each species but covered (less than ∼ 78,000 km^2^) in disaggregated pixels; however, main reduction areas were in the Asia (southern China and Mongolia). The broadest range expansions for *L. agraria* were anticipated in Asia (China, Russia, and Mongolia). In contrast, for *L. ingenua*, our results did not show a homogeneous pattern of potential range expansion; however, we noted increases in suitability in the Americas, Africa, Asia, Europe, and Australia. The biggest changes in distributional potential of *L. sativae* were in North America and western parts of South America (Figure 1). New potential range areas were also in Alaska and Canada (Figure 1). *Lycoriella agraria* and *L. sativae* potential range overlap was indicated in the western United States (Nevada, Arizona, Idaho, Wyoming, and Colorado) (Figures 1, and 2). Potential range overlap of *L. agraria* with *L. ingenua*, and *L. ingenua* with *L. sativae* were noted in central and western China (Qinghai, Xizang, and Xinjiang), central Kazakhstan, northern and northwestern Mongolia, northern Siberia, and the border regions between China and Mongolia (Figures 1, and 2).

## 4. Discussion

It is generally accepted that environmental changes will modify species’ geographic distributions worldwide [39]. Understanding how these changes will influence species’ distributions is particularly key for economically important species. The Sciaridae occurs almost worldwide [10], including important pests in mushroom crops, for example, [3], mainly in the genera *Bradysia* and *Lycoriella* [6].

*Lycoriella* includes the most threatening pests (e.g. our three species), causing important damage to mushroom production [4]. In Korea, the most economically important oyster mushroom pest is *L. ingenua*, among the six mushroom fly species [11]. Usually, *L. sativae* is the most abundant in fields, but is much less damaging than *L. ingenua* in mushroom culture [3].

How climate change will affect the geographic distributions of economically important sciarid species remains an open question. According to Sawangproh et al. (2016), ambient temperature can affect not only the survival and larval development of sciarid flies but also their feeding activity. As such, damage in mushroom crops or nurseries will be influenced by lower or higher temperatures. Apart from regional species checklists, little is known about the factors that drive these species’ distributions, so consequently little is known about impacts of climate change on the future distributions of these species. These insects are easily transported by human activities and, once they reach a suitable environment, they can build up populations, which can lead to major economic losses and establish populations in mushroom production areas.

Few studies have investigated the presence of sciarids in the Afrotropical region. Chidziya et al. (2013) considered *L. ingenua* (as *L. mali*) as the most damaging mushroom fly in Zimbabwe, but provided no occurrence records for the species. Katumanyane et al. (2020) reported for the first time the presence of both *L. ingenua* and *L. sativae* in South Africa. Our model has predicted suitable environmental conditions for these species in the southern portion of the African continent, including the above-mentioned countries (Figures 1, and 2), though no points from either country were included in the dataset used in model calibration.

The dominant and most serious pest species in mushroom crops in North America is *L. ingenua* [12]. Our results show that, for the USA, for example, current environmental suitability for this species is moderate for the entire West Coast and most of the southeastern part of the country, including most of the East Coast (Figure 1 and supp. information figure 5). Most of California presents high environmental suitability for the species, which is particularly relevant because California ranks second in the number of mushroom growers in the country, following only Pennsylvania [9].

Pennsylvania itself has moderate current environmental suitability (Figure 2), and our model predicts stable environmental suitability for the state under future scenarios (supp. information figure 5). These results should be taken into consideration, since it could lead to major economic losses to mushroom producers, considering that about 66% of all US mushroom growers are located in this state [9].

In South America, on the other hand, mushroom production is still incipient. It plays a growing social role as it becomes a different source of income for producers at local level. Brazil is the most outstanding case in South America, although efforts to cultivate mushrooms are beginning in other countries [43].

So far, no official record of species of *Lycoriella* exists for Brazil. Our model showed high environmental suitability in most of southern and southwestern Brazil for *L. ingenua* and *L. sativae* (Figure 2). As such, once these species are introduced in the country, they will likely have the ability to establish stable populations, a fact that must be regarded with caution because most Brazilian mushroom production is concentrated in the southern and southwestern states. Introduction of *Lycoriella* species to the country would pose an extra threat for Brazilian mushroom growers, who already face problems with other sciarid and scatopsid species [44,45].

The genus *Lycoriella* significantly reduces mushroom production inside greenhouses; these species also may impact other agricultural species (e.g. strawberry, nursery plants [6,46,47]. Our results show areas with suitable conditions for these flies around the world (Figure 2). We are particularly concerned about greenhouse availability, although we are not incorporating possible competition with other species in our models. However, *Lycoriella* species show very broad ecological niches with high possibilities invasive potential, from Brazil to Alaska. We suggest that experimental physiological studies that address the fundamental niche of these species more directly will be an important next step in protecting food production in greenhouses, to characterize areas with environmental conditions that characterize the physiological limits adequate to the development of *Lycoriella* populations.

## Authors’ contributions

**RM:** Conceptualization, Analysis, Writing Original Draft, Supervision, Project Administration, Analysis and Construction.

**JD**: Conceptualization, Data Curation, Writing Original Draft, Discussion.

**RK**: Conceptualization, Discussion.

**AF**: Writing Original Draft, Data Curation, Discussion

**CU**: Writing Original Draft

**DJG**: Conceptualization, Analysis, Writing Original Draft, Discussion.

## 5. Acknowledgments

We thank A. Townsend Peterson for comments and suggestions. We thank the Coordenação de Aperfeiçoamento de Pessoal de Nível Superior (CAPES) for the Ph.D scholarship awarded to Roberta Marques, for financial support in the “sandwich” program (Process: 47/2017), and for grants awarded to the second and third authors (CAPES grant numbers 88882.306693/2018-01 and 88882.182253/2018-01, respectively).

